# Breaking free from references: a consensus-based approach for community profiling with long amplicon nanopore data

**DOI:** 10.1101/2024.07.04.602031

**Authors:** Willem Stock, Coralie Rousseau, Glen Dierickx, Sofie D’hondt, Luz Amadei Martínez, Simon M Dittami, Luna van der Loos, Olivier De Clerck

**Affiliations:** Phycology Research Group, Ghent University, Belgium; Sorbonne University, CNRS, Integrative Biology of Marine Models (LBI2M, UMR 8227), Station Biologique de Roscoff (SBR), 29680 Roscoff, France; Research Group Mycology, Ghent university, Belgium; Research Institute for Nature and Forest, Geraardsbergen, Belgium; Laboratory of Protistology and Aquatic Ecology, Ghent University, Belgium

**Keywords:** Oxford Nanopore Technology, amplicon sequencing, chimera, consensus sequence

## Abstract

Third-generation sequencing platforms, such as Oxford Nanopore Technology (ONT), have made it possible to characterise communities through the sequencing of long amplicons. Whilst this theoretically allows for an increased taxonomic resolution compared to short-read sequencing platforms such as Illumina, the high error rate remains problematic to accurately identify the community members present within a sample. Here we present and validate CONCOMPRA, a tool that allows the detection of closely related strains within a community by drafting and mapping to consensus sequences. We show that CONCOMPRA outperforms several other tools for profiling bacterial communities using full-length 16S rRNA gene sequencing. Since CONCOMPRA does not rely on a sequence database for profiling communities, it is applicable to systems and amplicons for which a reference framework is poorly developed. Our validation test shows that the amplification of long PCR products is likely to produce chimeric byproducts that inflate alpha diversity and skew community structure, stressing the importance of chimera detection. CONCOMPRA is available on GitHub (https://github.com/willem-stock/CONCOMPRA).

**Key points:**

- We introduce a reference free tool for (long) amplicon Oxford Nanopore sequencing data
- We show that it is capable of outperforming achieving a higher accuracy than preexisting tools
- Chimera detection and removal is an essential step in the processing of long amplicon Oxford Nanopore sequencing data

## Background

Oxford Nanopore Technology (ONT) allows for much longer amplicons to be sequenced than second-generation sequencing platforms. Various studies have tried leveraging the increased length of marker amplicons to increase the taxonomic resolution at which microbial communities can be characterised. For instance, D’Andreano et al. (2021) sequenced a 3.5 Kb and a 6 Kb region, covering multiple rRNA genes and internal transcribed spacers regions, to classify fungi in clinical samples. van der Loos et al. (2021) described a protocol to characterize the bacterial communities associated with seaweeds by sequencing the near full-length 16S rRNA gene.

Despite continuous improvements in sequencing chemistry, flow cells and base calling algorithms, the quality of sequencing data is still markedly lower than that generated by second-generation short-read sequencing platforms. Due to the relatively high number of erroneous bases (modal read accuracy for the R9.4.1 and R10.4 flow cell of >96% and >99%;Luo et al., 2022), the increased amplicon length obtained with ONT sequencing poses challenges in itself and does not necessarily result in a finer phylogenetic resolution. The direct taxonomic assignment of the full-length 16S rRNA gene at the single-read level, using established tools such as EPI2ME platform is for instance problematic at subgenus level (Winand et al., 2019). One way to increase read accuracy is to ensure that the same DNA string and its complement or copies, are sequenced multiple times. This can be achieved with unique molecular identifiers that link daughter sequences to the original template DNA (Karst et al., 2021) or rolling circle amplification (Baloğlu et al., 2021; Calus et al., 2018). The downside from such approaches is that they complicate the workflow, raise costs and reduce the number of unique reads that can be sequenced within a run.

Given these complications, researchers have focused on developing post-sequencing solutions to obtain species level resolution with ONT amplicon data. Emu (Curry et al., 2022)employs an expectation-maximization algorithm to provide species-level taxonomic assignments from full-length 16S rRNA reads. NanoCLUST classifies consensus sequences instead of the individual reads (Rodríguez-Pérez et al., 2021). These tools, as well as other common tools used for taxonomic assignment of amplicon ONT sequencing data, including Kraken2 (Lu & Salzberg, 2020), require a reference database such as SILVA (Yilmaz et al., 2014), UNITE (Abarenkov et al., 2024) or PR2 (Guillou et al., 2013) to function. The databases available to date, however, are far from exhaustive and often lack strain or even species-level information, hampering finer taxonomic assignment of long amplicon ONT sequencing with the available tools. This is, for instance, the case for marine bacterioplankton, since many of these oligotrophic bacteria are challenging to isolate (Henson et al., 2016; Katayama et al., 2024). Moreover, databases are only available for a few loci (ITS, rRNA genes), and might not be suitable when working on an understudied group or in unexplored habitats.

As with other sequencing technologies, ONT generated sequences can be chimeric. Chimeras in amplicon sequencing data can easily make up one third of the data (C-y Wang & Wang, 1997) although the frequency of chimeric reads will depend on the sequencing protocol (i.e. primers used, PCR conditions) as well as the samples composition itself (Ho et al., 2021; Qin et al., 2023). Chimeras are formed during the PCR as well as post-amplification, during the ligation (Eccles et al., 2017). The amplification of a longer region generally produces many more byproducts (Laver et al., 2016) as template switching during DNA synthesis is more likely to occur. Several tools have been developed to detect and remove chimeric reads in ONT sequencing data, e.g. Liger2LiGer (Marijon et al., 2020, https://github.com/rlorigro/Liger2LiGer), but these are not designed to handle amplicon data and are computationally demanding as they require all-against-all mapping of reads. Mapping of the reads with chimeric sequences still present is likely to result in false positive detection of species and inflate diversity estimates.

We present the tool CONCOMPRA, CONsensus approach for COMmunity Profiling with nanopoRe Amplicon sequencing data. This tool creates a consensus sequence database followed by abundance profiling of amplicon ONT sequencing data. Our workflow is intended to be fast, customizable (implemented in bash and python) and flexible (working from one-to-many samples). We show that this workflow, using full-length 16S rRNA gene data, allows for distinguishing closely related species and works with long amplicons for which no suitable reference database is available.

## Methods

### CONCOMPRA workflow

The input data consist of base-called ONT sequencing reads in fastq format. CONCOMPRA (Figure 1) will process all fastq files present in the appointed working directory. On a file per file basis, a list of consensus sequences will be generated from the reads according to the following steps. Reads outside of the user-defined length window are discarded. Forward reads with primers in the expected positions are identified using primer-chop (https://gitlab.com/mcfrith/primer-chop), which also trims the primers and bases preceding the forward primer and trailing the reverse primer. Simultaneously, the mapping of the primers to the sequences is used to estimate the rates (probabilities) of insertion, deletion, and substitutions in the sequencing data (Hamada et al., 2017), required for LAST alignment algorithm (Kiełbasa et al., 2011) implemented in primer-chop and our consensus generation approach. The top 80% of the forward, trimmed reads, with the fewest expected errors is retained using Filtlong (https://github.com/rrwick/Filtlong). These sequences are used for unsupervised clustering, similar to the NanoCLUST approach. The reads are clustered with UMAP-OPTICS (Malzer & Baum, 2020; McInnes et al., 2018) based on 3mer composition. A consensus sequence is produced from a maximum of 40 randomly selected sequences in each cluster using lamassemble (Frith et al., 2021) with the previously estimated insertion, deletion, and substitution rates. Potentially chimeric sequences are flagged using the vsearch uchime_denovo algorithm, considering the number of reads assigned to the cluster from which the consensus sequence was drafted.

**Figure 1:**
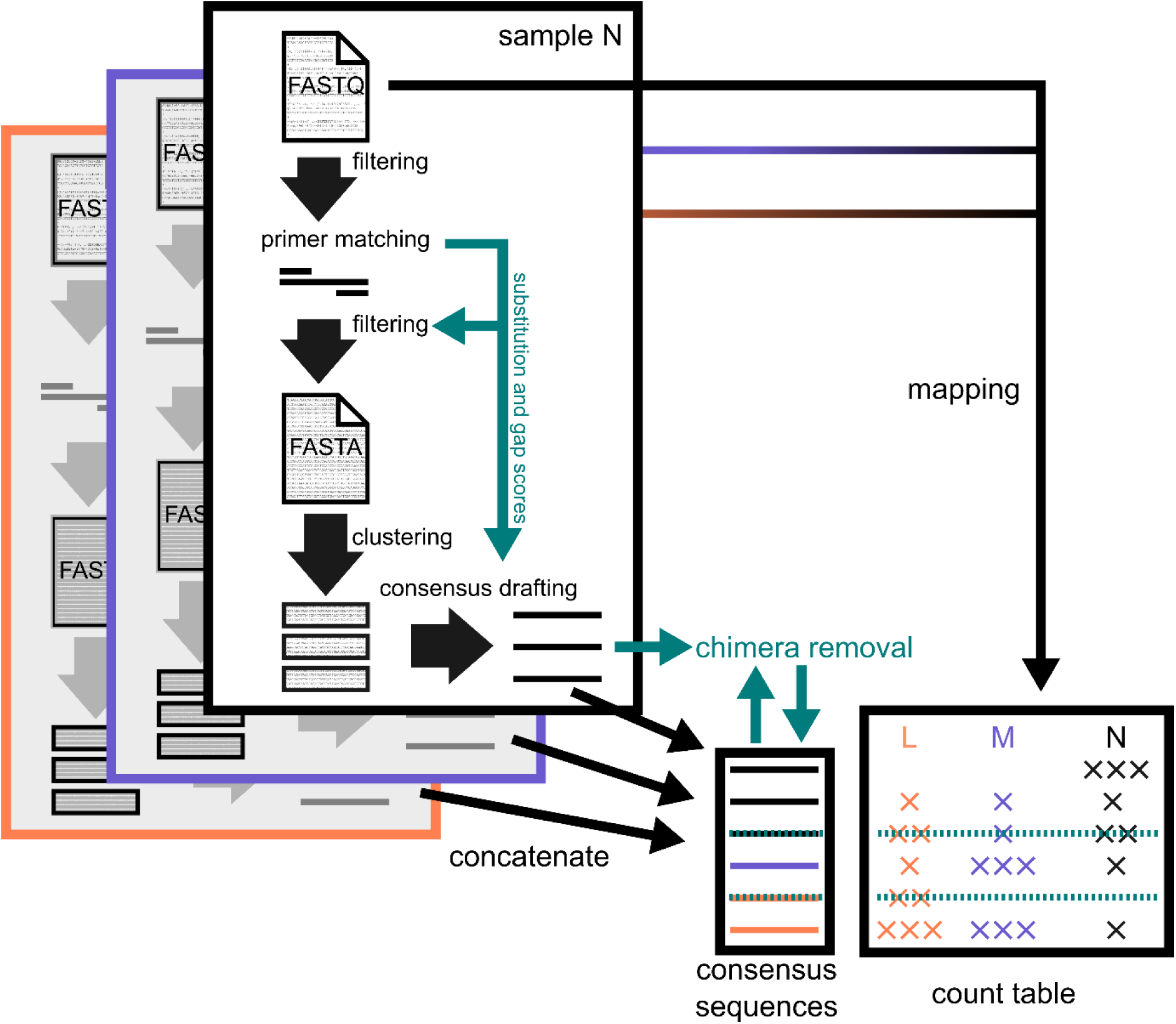
CONCOMPRA workflow. Consensus sequences are drafted for every sample individually and then concatenated across samples. Sequences are mapped to the consensus sequence table to generate a count table with the number of sequences mapped to each consensus sequence per sample. Chimeric sequences are detected and removed.

Vsearch (Rognes et al., 2016) is used to deduplicate sequences and, optionally, remove highly similar consensus sequences coming from the different samples. An abundance table across samples of the consensus sequence is created by mapping the reads from the original fastq files to the consensus sequences with minimap2 (Li, 2018). Reads that fail to map to the consensus sequences are written to a separate ‘unmapped’ fastq file. Sequences that were flagged as chimeric during the per sample drafting of the consensus sequences as well as the sequences that are deemed chimeric based on the number of reads mapped to the consensus sequences across samples (vsearch uchime_denovo) are removed in order to obtain a final, chimera free consensus sequence table.

### Datasets

CONCOMPRA was compared to alternative tools on two datasets: 1) the 16S rRNA gene sequences from a synthetic bacterial community (16S MOCK), where the ground truth was known, and 2) low-quality 16S rRNA gene sequences from natural planktonic bacterial communities (16S natural).

16S MOCK: The full-length 16S rRNA gene of 1 µl of a synthetic bacterial community, composed of evenly mixed genome DNA of 20 bacterial strains, (ATCC MSA-1002™), was amplified with the primer set 27F_Bctail-FW (TTTCTGTTGGTGCTGATATTGC_AGAGTTTGATCMTGGCTCAG) and 1492R_Bctail-RV (ACTTGCCTGTCGCTCTATCTTC_CGGTTACCTTGTTACGACTT) using the Phire Tissue direct PCR Master Mix (Thermo Fisher) as previously described (van der Loos et al. 2021). The PCR product was barcoded using the Oxford Nanopore PCR Barcoding Expansion Pack 1-96 (EXP-PBC096) and pooled equimolarly with various other samples. The pooled DNA was purified using the Agencourt AMPure XP system. The PCR and barcoding, starting from the synthetic bacterial community DNA extract was done four times independently, resulting in four different barcoded PCR products. The library was prepared for two different nanopore flow cell chemistries (two barcoded PCR products each), R9.4.1 and R10.4.1 flow cell, using respectively the ONT SQK*-*LSK109 and SQK-LSK114 (V14) ligations kits, with the Amplicon by Ligation protocol. Sequencing on the R9.4.1 flow cell was run for 48h and on the R10.4.1 flow cell for 72h. The neural network-based tool Guppy (Wick et al., 2019) was used for super-accurate basecalling (v6.3.8 for the R9.4.1 flow cell data; v6.5.7 for the R10.4.1 flow cell). The minimum data quality threshold was set to Q10. Data quality was assessed with NanoPlot v1.42.0 (De Coster & Rademakers, 2023). The first 10 000 reads of each sequence file were blasted (BLAST 2.2.29+) against the reference sequences (retrieved from the genomes provided by ATCC) to estimate sequence similarity of the data to the ground truth. An Illumina MiSeq system (300bp paired end) sequenced data set, restricted to the V1-V3 region of the same mock community, was generated using the pA (5′-TCGTCGGCAGCGTCAGATGTGTATAAGAGACAGAGAGTTTGATCCTGGCTCAG-3′) and BKL1 primer pair (5′-GTCTCGTGGGCTCGGAGATGTGTATAAGAGACAGGTATTACCGCGGCTGCTGGCA-3′) as in D’Hondt et al. (2018).

16S natural: Three independent water samples (50-100ml), collected in different months at different locations within the Belgian part of the Schelde river were filtered through a 0.22 µm MF-Millipore Membrane filter (Merck Millipore). DNA was extracted using Dneasy PowerLyzer Microbial Kit (Qiagen). For Illumina sequencing, the V1-V3 region of the 16S rRNA gene was amplified and sequenced as described above. The full-length 16S rRNA gene for the three communities was amplified and sequenced on a R9.4.1 flow cell as described for the synthetic bacterial community. Basecalling was performed using Guppy v 4.3.4. The resulting base-called data had less than 3% above the Q10 quality cutoff so the Q10 quality threshold was not enforced to prevent excessive data loss.

### Data analysis

CONCOMPRA was run on the datasets with sequence length window set to 1400-1700pb. The mock samples were analysed individually, and the natural samples were analysed in batch. Taxonomic annotations of the CONCOMPRA-created consensus sequences for the 16S MOCK and 16S natural dataset were obtained using the IdTaxa function from the DECIPHER package v2.28.0 (Murali et al., 2018)with the SILVA_SSU_r138 reference database (Quast et al., 2013). NGSpeciesID v0.3.0 (Sahlin et al., 2021)and amplicon_sorter (version Feb 20, 2024; Vierstraete & Braeckman, 2022) were used to compare quality of the obtained consensus sequences for both mock datasets. NGSpeciesID was run with an abundance ratio threshold of 0.002 and Racon as polisher using three iterations. Chopper v0.7.0 (De Coster & Rademakers, 2023) was used to retain only ≥Q12 reads prior to using amplicon_sorter. The same size window was set for amplicon_sorter as what was used for CONCOMPRA (1400-1700bp). Consensus sequences generated by CONCOMPRA, NGSpeciesID, and amplicon_sorter were compared to the reference sequences using BLAST 2.2.29+. Only reference with a >95% match to a consensus sequence were considered to be represented by the consensus sequences.

The 16S MOCK data and 16S natural data were also analyzed with Emu v3.4.5 and Kraken2 v2.1.2. SILVA_SSU_r138 was set as reference database. The 16S MOCK data was analysed by both reference-based tools as is, after length (1400-1700bp) filtering of the data with Chopper, and after scrubbing chimeric reads (i.e., removal of putative chimeric sequenced regions) with yacrd v1 (Marijon et al., 2020) on an all-to-all file generated by mm2-fast v2.24(Kalikar et al., 2022).

As scrubbing failed for one of the R10 datasets, this single sample was excluded for calculating statistics on filtering and scrubbing.

The Illumina data was processed with DADA2 (v1.28.0; Callahan et al., 2016). The primer regions as well as the trailing ends of low quality were trimmed prior to applying the default DADA2 workflow (v1.16). The IdTaxa function from the DECIPHER package v2.28.0 with the SILVA_SSU_r138 reference database was used for taxonomically annotating the sequence variants (SVs).

In order to compare the taxonomic annotations between approaches, taxonomic tables were aggregated at the genus level using Phyloseq (1.44.0; R 4.3.1; McMurdie & Holmes, 2013). Only bacterial reads were retained in the samples by removing reads assigned to mitochondria, chloroplasts or not the domain Bacteria. Reads that could not be annotated at the phylum level were also removed. Accuracy of the different methods was quantified by summing the absolute values of observed relative abundance minus the expected relative abundance across genera. In order to compare taxonomic resolution in the absence of a predefined taxonomic framework, the OTUs/SV assigned to the understudied NS11-12 marine group (Sphingobacteriales) within the 16S natural dataset were compared. An approximately-maximum-likelihood phylogenetic tree (FastTree 2.1.11, Price et al., 2010) was constructed from the OTU/SV sequences, which were aligned with SSU-ALIGN 0.1.1 (Nawrocki, 2009). *Sphingobacterium multivorum* OM-A8 (ENA accession id: AB020205) was used as outgroup for rooting.

## Results

### 16S mock community

The reads, generated by ONT sequencing (Table 1), had a median sequence similarity to the reference sequences of 92.0% and 94.4% for the R9 and R10 datasets respectively. The number of non-chimeric consensus sequences produced by CONCOMPRA was close to the number of bacteria present in the mock (19.5±2.4 sd) and the average sequence similarity of the consensus sequences to the best matching genome derived 16S rRNA gene sequences (https://www.atcc.org/products/msa-1002) was ≥99.9% for the four samples (Figure 2). Over half the reads were assigned to chimeric OTUs (59.3% ±4.5% sd), which is in line with the relatively low similarity of the reads to the actual reference sequences (Figure 2).

**Figure 2:**
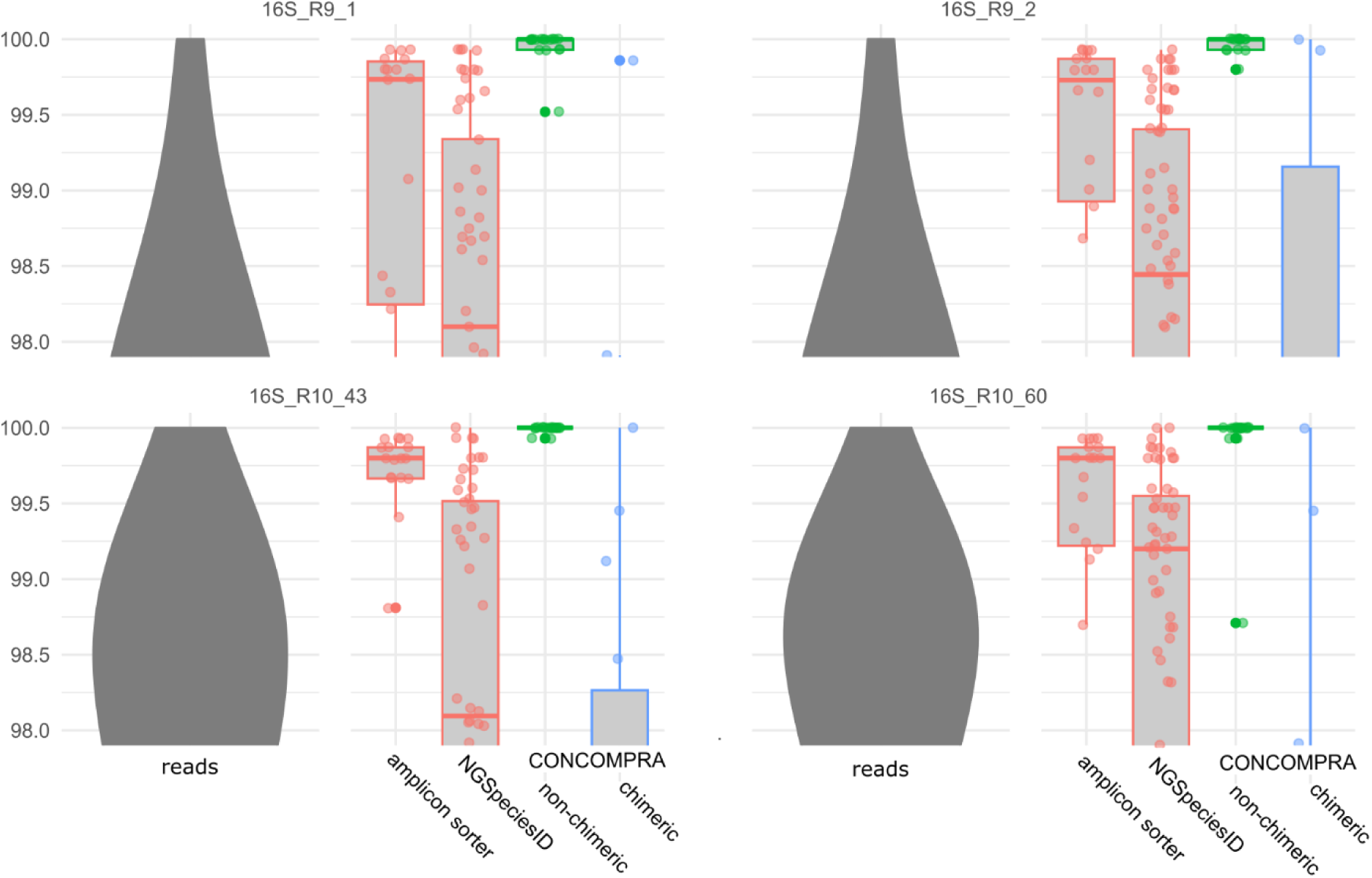
Similarity of the sequencing data and the consensus sequences to the reference 16S mock community. The blast-based percentage similarity to the references is shown for the different datasets (violin plots) and consensus sequences (box plots). The reads (dark grey) from the R10 sets (bottom row) were markedly more similar to the reference sequences (>20% of the reads were at least 98% similar to the references) than the reads from the R9 sets (>5% of the reads were at least 98% similar to the references; top row). The alternative consensus building tools (red) generated more consensus sequences, but these were less similar than the non-chimeric consensus sequences drafted by CONCOMPRA (green). The chimeric consensus sequences from CONCOMPRA were least similar (97% on average) to the reference sequences.

**Table 1:**
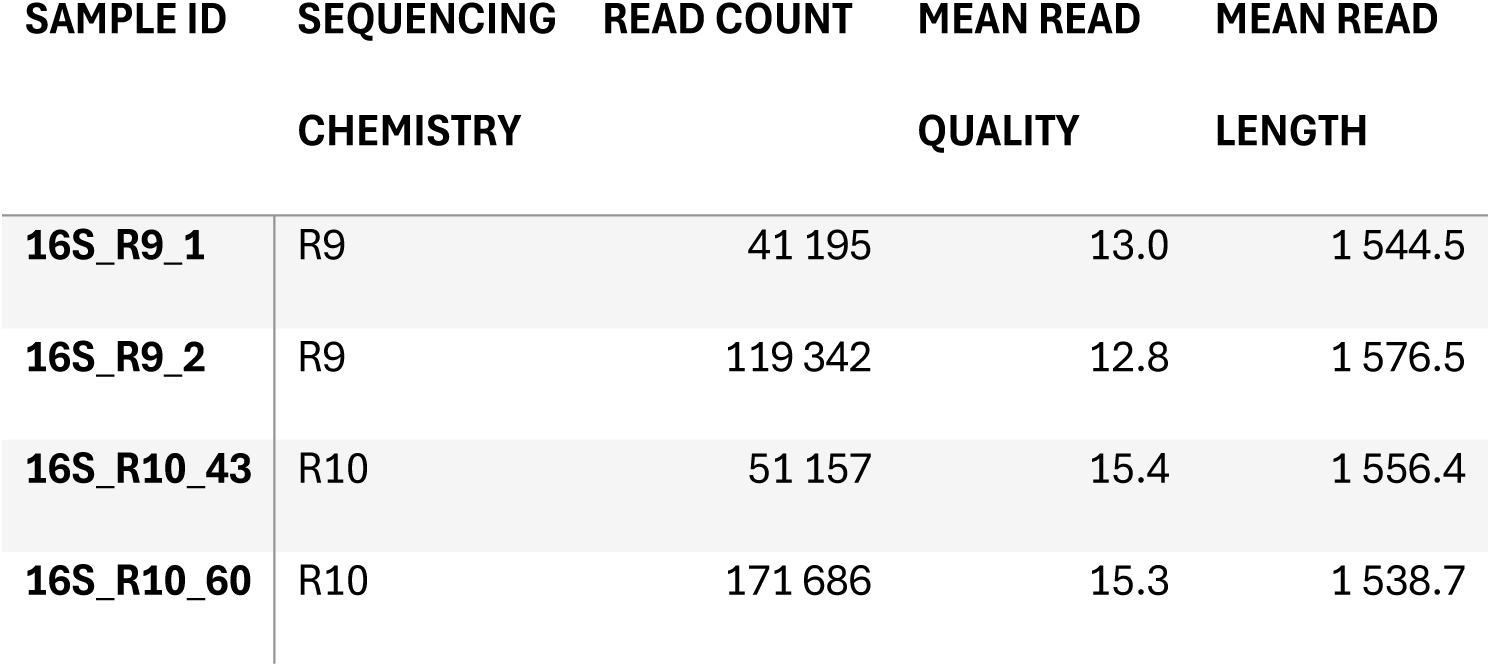
summary of the ONT 16S rRNA gene generated data for the mock community.

The consensus building tools NGSpeciesID and amplicon_sorter returned 77.3 (±12.1 sd) and 36.5 (±8.3 sd) consensus sequences respectively. Both tools were consistently less accurate than CONCOMPRA, obtaining an average sequence similarity of 99.1% (±0.3 sd) and 99,5% (±0.3 sd; Figure 2). CONCOMPRA was also the only tool that obtained consensus sequences that were identical to the reference sequences, which it did for over half the matching sequences (Figure 3).

**Figure 3:**
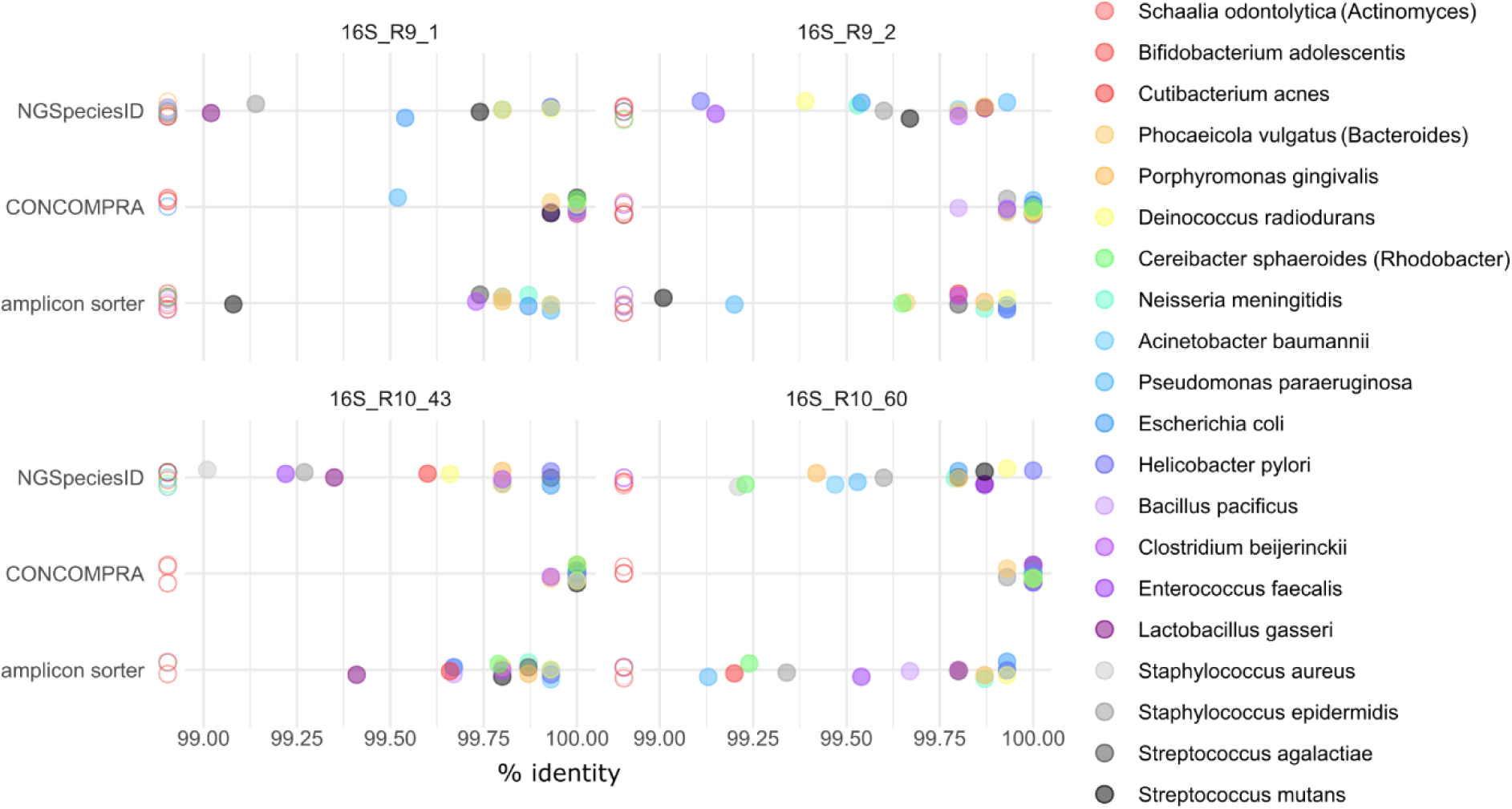
Detection of the 20 different species in the bacterial mock community. The highest similarity to any of the 16S rRNA gene copy sequences to each species (colours) is shown for the different consensus building tools (rows) for the different 16S datasets. Only non-chimeric sequences are considered for CONCOMPRA. Species for which no consensus sequences was present with an identify above 99% are shown as non-filled circles to the left of the axis.

No consensus sequences with a similarity above 95% were recovered for *Bifidobacterium adolescentis* or *Schaalia odontolytica* in any of the tests, suggesting failure to sufficiently amplify these 16S rRNA gene sequences (Figure 3). In the R10 datasets, *Cutibacterium acnes* was absent once in a CONCOMPRA consensus sequence set and *Bacillus pacificus* in a NGSpeciesID consensus sequence set. More bacteria remained undetected in the R9 datasets: *Cutibacterium acnes* was absent in both CONCOMPRA sets and *Escherichia coli*, *Enterococcus faecalis* and *Streptococcus agalactiae* were each absent once in the CONCOMPRA sets. *Bacillus pacificus* was missing in both amplicon_sorter sets and *Cereibacter sphaeroides* in one NGSpeciesID set. The two *Streptococcus* strains were distinguished by all tools. CONCOMPRA was the only tool that could consistently distinguish the *Staphylococcus* strains, although the consensus sequences for the *S. agalactiae* in one of the R9 sets was only 95% identical to the reference sequence.

The two alternative, reference-based community profiling tools returned a higher number of OTUs for the mock communities than COMCOMPRA. Kraken2 resulted in 500 (±78.7 sd) OTUs, and Emu predicted 80.7 (±14.2 sd) OTUs. Filtering out the sequences outside the size window, removing about 16% of the sequences, reduced the predicted number of OTUs (Kraken2: 339.3±48.3sd; Emu: 68±11.4sd). Scrubbing the reads, removing 0.2% of the reads, did not reduce the number of predicted OTUs (Kraken2: 505±76.6sd; Emu: 83.3±13.4sd). Community profiles remained highly similar, independent of data treatment (Figure 4). The Illumina data, processed by the DADA2 pipeline, returned 185 sequence variants (SVs) and was distinct from any of the ONT processed datasets (Figure 4). The Illumina data were highly skewed: *Clostridium* making up over half the community, whilst *Streptococcus* and *Staphyloccoccus* were present at less than 1%. CONCOMPRA approximated the real abundance profile best (summed deviation CONCOMPRA = 46.9 ± 6.2% sd; Kraken2 = 52.7± 1.6% sd; Emu = 55.7± 1.7% sd; Illumina = 139.1). Filtering out the sequences outside of the set size window did not reduce the deviation of the two alternative community profiling tools (Kraken2 = 53.6± 2.5% sd; Emu = 55.7± 1.4% sd).

**Figure 4:**
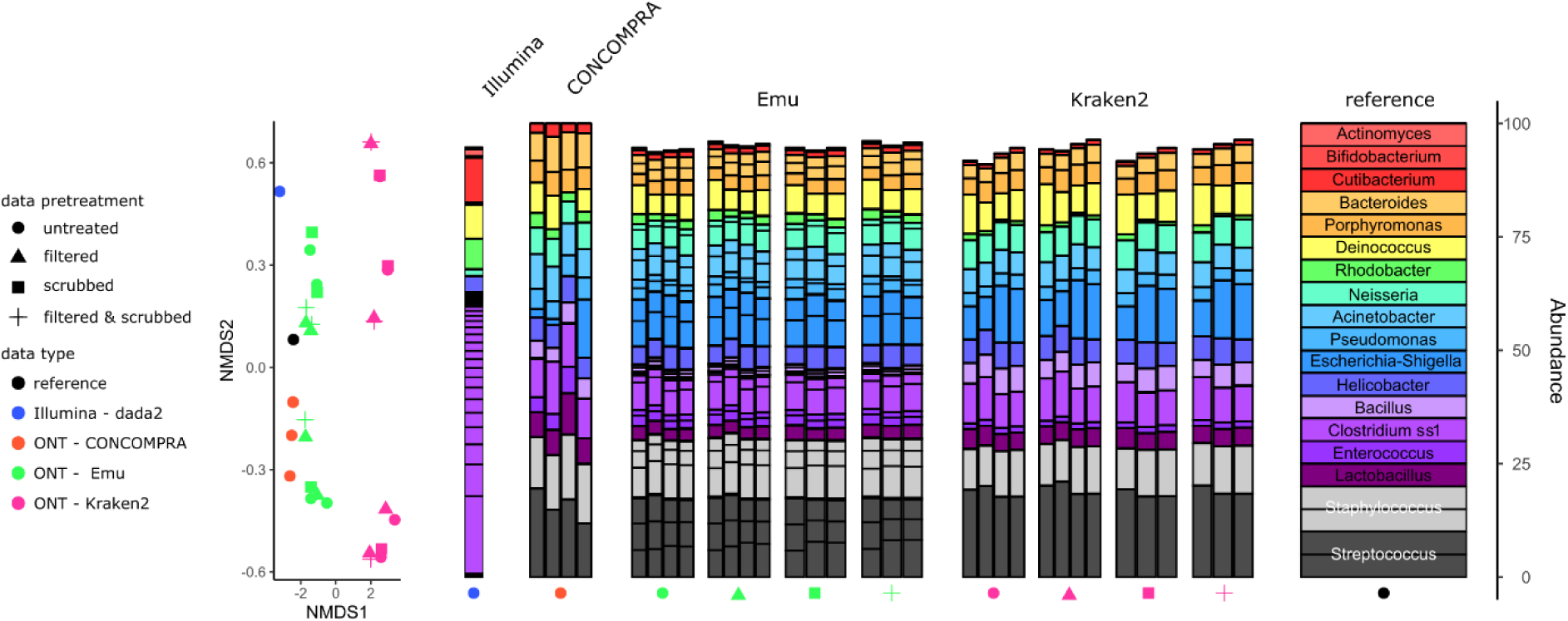
inferred composition of the mock communities. The beta diversity is visualised through non-metric multidimensional scaling (NMDS; left) using Bray-Curtis dissimilarity on the relative abundances of the genera (2D stress = 0.037). Colours are used to indicate the data types and pipelines used and the shape reflects the data treatment. The community composition (right) differed most between the Illumina data and the ONT data. Only the genera present in the reference community are shown in the bar plots. Nomenclature from the SILVA_SSU_r138 referencereference database is used.

### Natural bacterial samples

CONCOMPRA identified 64 nonchimeric OTUs across the three samples, assigned to 15 different bacterial genera. Kraken2 assigned the ONT reads to 1894 different bacterial genera (2585 taxa). This was substantially less with Emu, namely 265 different bacterial genera (506 taxa). Using the pair-end Illumina data, DADA2 identified 948 SVs assigned to 81 different bacterial genera. Although CONCOMPRA identified the fewest genera out of the three ONT based workflows, 81.7 ± 4.7% (sd) of the assigned reads in CONCOMPRA were allocated to genera also found in the Illumina based DADA2 dataset. This was far less for the reads within Kraken2 and Emu (Kraken2: 16.6± 2.6%; Emu: 40.6 ± 7.3%). The reverse is also true as the majority (67.4 ± 10.7%) of the reads from the DADA2 datasets were assigned to the genera found by CONCOMPRA.

The Illumina-DADA2 and CONCOMPRA captured a similar trend in community composition (Figure 5), although differences between communities were more pronounced for Illumina-DADA2 datasets. The compositional accumulated difference between Illumina data and ONT based approaches were largest for Kraken2 (Kraken2: 116.7 ± 7.6% sd; Emu: 83.0 ± 6.9%; CONCOMPRA 81.7 ± 17.6% sd).

**Figure 5:**
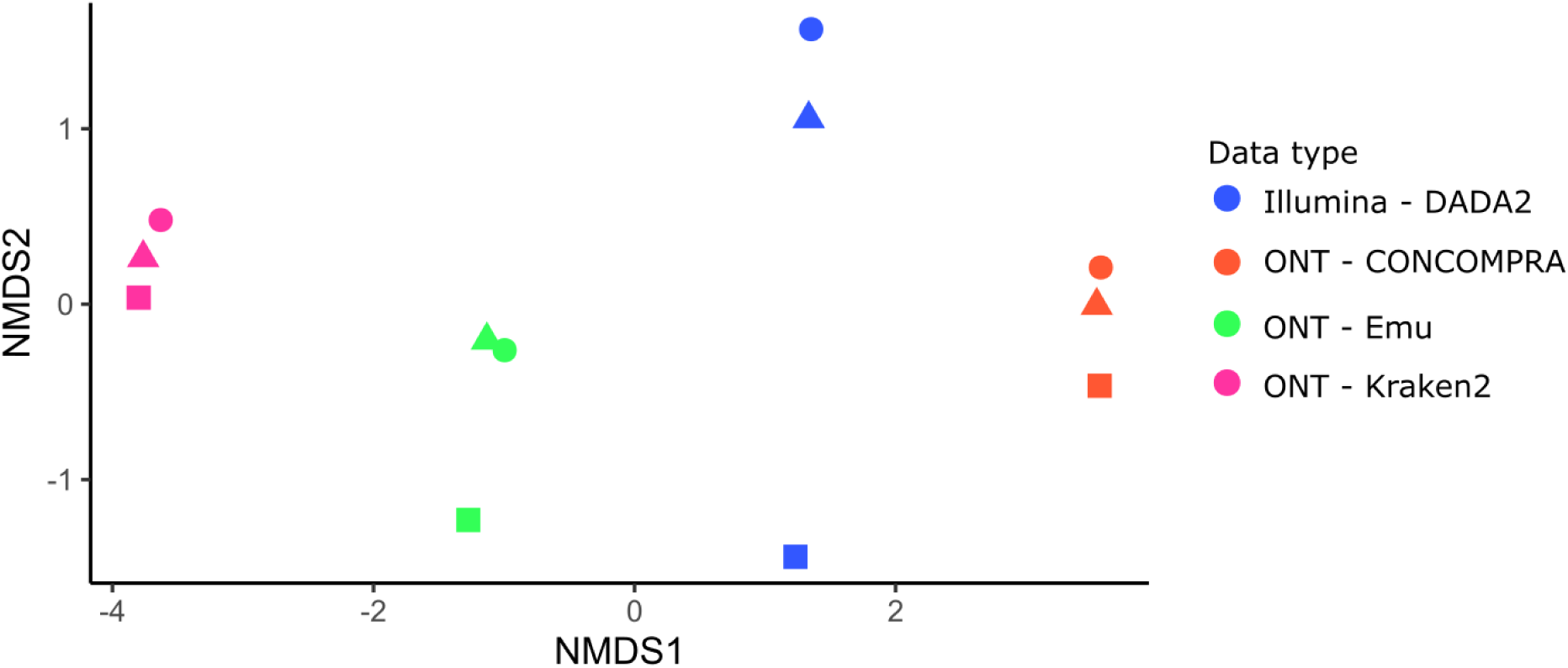
The differences in community composition between natural samples are most pronounced based on Illumina sequencing but the overall trends are conserved across sequences approaches and analyses. The beta diversity is visualised through non-metric multidimensional scaling (NMDS) using Bray-Curtis dissimilarity. Symbols are used to indicate the different natural bacterial communities and colours are used to represent the different analyses types. Stress = 0.04

The NS11-12 marine group (Sphingobacteriales) was detected in all three samples, with all methods (Figure 6), although their relative abundance differed between the methods. Kraken2 and Emu datasets only had a single taxonomic ID affiliated with the NS11-12 marine group, whilst CONCOMPRA and DADA2, respectively, drafted four and six different SVs assigned to this group. These different sequences could be assigned to two different clades (Figure 6). CONCOMPRA identified predominantly members from the first NS11-12 clade in the first two samples and from the second clade in the third sample. This is in line with the output from DADA2.

**Figure 6:**
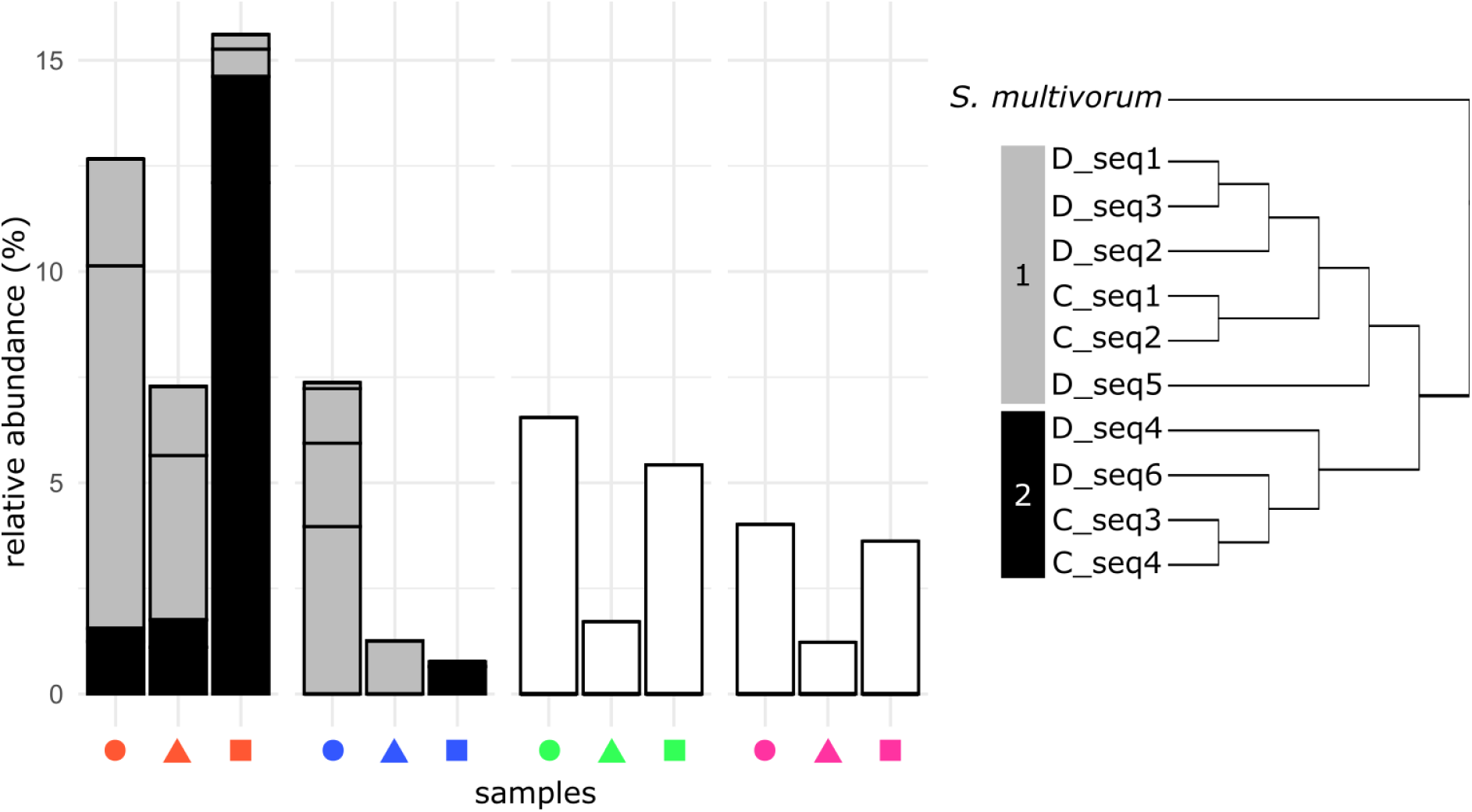
the relative abundance of the NS11-12 marine group in the natural 16S samples. The different clades within the NS11-12 marine group (right) are indicated in the bar plots by different colours. The sequences from the DADA2 and CONCOMPRA dataset are indicated in the phylogenetic tree with a ‘D’ and a ‘C’ in the tip labels, respectively.

## Discussion

We presented the results from a novel tool that, starting from ONT amplicon sequencing data, creates a *de novo*, consensus-based sequence database which is then used for abundance profiling of microbial communities. This tool, CONCOMPRA, proved to be more sensitive and generally more precise than the other consensus sequence generating tools tested here. CONCOMPRA performed well in samples with a low (20 strains) diversity and was able to capture the diversity of abundant taxa in natural samples.

In natural, more diverse samples, the bacterial diversity reported by CONCOMPRA was lower than the output returned by Emu and substantially lower than the diversity results from Kraken2. Based on the diversity reported by DADA2, the real bacterial diversity within those samples, at least in terms of number of genera present, is likely in between the CONCOMPRA and Emu results. As both CONCOMPRA and Emu resulted in datasets which were compositionally similar to the DADA2 datasets, the differences between CONCOMPRA and Emu reflect the trade-off between sensitivity and specificity.

Reference databases, even of established marker genes such as the 16S rRNA gene that was used in this study, are incomplete. This is the case for the NS11-12 marine group, Bacteroidota which are important in structuring the coastal and estuarine planktonic bacterial community (Korlević et al., 2022; Wang et al., 2021). As this group is only represented in the SILVA_SSU_r138 database by a single taxon ID, Kraken2 and Emu could not distinguish the different clades present in the natural samples. CONCOMPRA on the other hand, was able to assign the reads to two clades within the NS11-12 marine group, in line with the output from the DADA2 workflow. This example illustrates well the benefit of being independent from an existing reference database for assigning reads to OTUs.

Our results showed that the bulk of the ONT sequencing data was less similar to references than can be expected based on reported base calling error rates (Luo et al., 2022). This is in line with the expected high frequency of chimeric reads in the data. The prevalence of chimeras hampered tools like NGSpeciesID to draft accurate consensus sequences and inflated the estimated richness by the Emu and Kraken2. This warrants the inclusion of extensive chimera detection and removal steps in long-read amplicon sequencing data workflows. Whilst yacrd, which was developed for chimera removal genome assembly rather than handling amplicon sequencing data (Marijon et al., 2020), failed to remove most chimeras, the vsearch uchime_denovo algorithm, as implemented in CONCOMPRA, removed a substantial amount of likely chimeric reads. By only removing chimeric consensus sequences after mapping the reads in our workflow, we aimed to increase mapping accuracy. This approach is only feasible in a workflow that relies on drafting de novo consensus sequences. The use of a de novo drafted reference database, as is also the case in most Illumina processing workflows (Callahan et al., 2016; Estaki et al., 2020; Rognes et al., 2016) can therefore be considered an asset over mapping to existing reference database as is now implemented in most Kraken2 or Emu workflows. Modifications to the sequencing protocol such as the optimalisation of PCR conditions (Fujiyoshi et al., 2020), are likely to reduce the number of chimeric reads and are important to consider as well.

Many workflows have been developed to process ONT sequenced (long) amplicon data, most of which rely on mapping reads to existing databases (Ammer-Herrmenau et al., 2021; Bertolo et al., 2024; Curry et al., 2022; Lu & Salzberg, 2020). Here we argued that there are important benefits to breaking free from reference databases and using a de novo approach instead. We introduced and validated a novel tool, CONCOMPRA, that drafts and maps to consensus sequences, thus allowing the distinction of closely related strains without any external reference database. It remains challenging to deal with the relatively high base calling error rates and common chimeric reads inherent to long amplicon Nanopore sequencing data. Overcoming these nontrivial issues is an essential step to maximally take advantage of the long-amplicon capability specific to the ONT platforms.

## Data availability

The sequencing data from the synthetic bacterial communities analyzed in this article is available in the NCBI BioProject database at https://www.ncbi.nlm.nih.gov/bioproject/, and can be accessed with PRJNA1129458. CONCOMPRA is available on GitHub (https://github.com/willem-stock/CONCOMPRA)

## Ethics statement

This research has been conducted in a fair and ethical manner, according to international and local guidelines. No human or animal derived data or test subjects were involved.

